# Investigation of biochemical changes in barley inoculated with *Trichoderma harzianum* T-22 under salt stress

**DOI:** 10.1101/2020.10.19.346056

**Authors:** Sneha Gupta, Penelope M. C. Smith, Berin A. Boughton, Thusitha W. T. Rupasinghe, Siria H.A. Natera, Ute Roessner

## Abstract

Increases in soil salinity impacts growth and yield of agricultural plants by inhibiting plant functions. Soil salinization is increasing because of the pressure of a growing population on food supply. Genetically modified crops and plant breeding techniques are being used to produce plants tolerant to salt stress. However, interactions of fungal endophytes with crop plants can also improve tolerance and is a less expensive approach. Here, the role of *Trichoderma harzianum* T-22 in alleviating NaCl-induced stress in two barley genotypes (cv. Vlamingh and cv. Gairdner) has been investigated. Metabolomics using GC-MS for polar metabolites and LC-MS for lipids was employed to provide insights into the biochemical changes in barley roots inoculated with fungus during the early stages of interaction. *T. harzianum* increased the root length of both genotypes under controlled and saline conditions. The fungus reduced the relative concentration of sugars in both genotypes and caused no change in organic acids under saline conditions. Amino acids decreased only in cv. Gairdner in fungus-inoculated roots under saline conditions. Lipid analyses suggest that salt stress causes large changes in the lipid profile of roots but that inoculation with fungus greatly reduces the extent of these changes. By studying a tolerant and a sensitive genotype and their responses to salt and inoculation we have been able to develop hypotheses about what lipid species and metabolites may be involved in the tolerant genotype for its tolerance to salt and how fungal inoculation changes the response of the sensitive genotype to improve its tolerance.

## Introduction

Soil salinity is becoming a significant problem worldwide as it is encountered in all climates (Shahbaz and Ashraf, 2013). We need to understand the crop response to salinity to minimize economic loss and improve crop productivity. The negative effects of salinity on growth, biomass and grain yield, and in causing oxidative damage, are well documented for several crops (Shahbaz and Ashraf, 2013). The extent of the damage caused by salinity depends on the plant’s developmental stage, duration of the salt stress, the concentration of salt, and the genotype (Hu et al., 2017). At the plant organ level, shoots are more sensitive to salinity than roots. However, roots are first exposed to salinity and respond to it rapidly.

The study of root growth has become more prevalent over a period of time amongst biologists (Yu et al., 2020). Among all possible root traits that can be measured, specific root length (SRL) and root length density (RLD) are of particular interest. However, one must know the total root length (TRL) in the plant before calculating SRL and RLD (Delory et al., 2017). This trait of measuring TRL has been used to measure the effect of salt stress on cereal crops such as rice (Kakar et al., 2019), wheat (Rahnama et al., 2019) and barley (Sarabia et al., 2019).

Roots are crucial for a myriad of physiological processes including water and nutrient uptake, anchoring plants and providing mechanical support to aboveground organs. Roots also affect surrounding nutrient composition by release of organic compounds that play a vital role in mineralizing nutrients (de Willigen and van Noordwijk, 1987). The area surrounding the root system, the rhizosphere, not only consists of root tissues but is home to many microorganisms that influence plant functions. Salt stress influences the root system (Yu et al., 2020) which respond by secreting compounds to protect against negative effects and encourage positive microbial interactions (Munns and Tester, 2008). These secreted compounds usually induce an interactive metabolic crosstalk involving diverse biosynthetic networks and pathways. Tolerance to salt stress involves dynamic changes in gene expression and metabolic pathways resulting in physiological adaptations (Gregory, 2006). Mitigation strategies that allow plants to deal with salt stress include osmolyte accumulation, activation of antioxidant enzymes and synthesis of compounds to combat reactive oxygen species (ROS), and modulation of hormones (Hossain and Dietz, 2016). Interaction with salt tolerant microbial inoculants can also enhance crop growth in saline soils (Morgan et al., 2005). Microbes that interact with plants have a diverse array of metabolic and genetic strategies that reduce the impact of salt stress and other abiotic stresses arising from extreme environmental conditions (Meena et al., 2017).

Significant progress has been made in the past decade to understand the mechanisms of salt tolerance imparted by endophytic fungi. *Trichoderma* species are one group with beneficial effects on plants under saline conditions. *T. virens* promoted Arabidopsis growth through the auxin response pathway to modulate root system architecture and activate auxin regulated gene expression (Contreras-Cornejo et al., 2009). Arabidopsis seedlings inoculated with *T. atroviride* developed a more branched root system than uninoculated plants, had enhanced shoot growth and increased chlorophyll content, demonstrating the beneficial effects of fungus under salt stress (Contreras-Cornejo et al., 2014). *T. harzianum* strain T22 and *T. longibrachiatum* T6 increased root length in tomato (Mastouri et al., 2010) and wheat plants (Zhang et al., 2016), when compared to uninoculated plants under saline conditions.

Colonization of several crops by endophytic fungi mitigates stress by increasing the levels of protective metabolites and osmoprotectants, activating antioxidant systems to prevent ROS damage, and modulating phytohormones to minimize salt effects (McNear Jr, 2013). Most efforts in characterizing and defining the biochemical changes related to the response during plant-microbe interactions have been driven by measuring any one class of metabolites (Tugizimana et al., 2018). However, there are still gaps in holistically understanding the dynamics and complexity of molecular mechanisms involved in the interactions, mainly due to the complexity of the multi-layered biological information networks involved (Tugizimana et al., 2018). Although the effects on plant biomass of the interactions are well described, the metabolic contributions of endophytes to host plants is less well documented. The early stage of the interaction is especially crucial in priming plants to respond to the environment. The alteration of plant metabolic responses during this stage has been suggested to correspond to the environment in which the interactions occur (Grayson, 2013).

In this study we investigated the interaction of two barley genotypes (cv. Vlamingh and cv. Gairdner) with contrasting germination phenology and salinity tolerance inoculated with the beneficial endophytic fungi *T. harzianum* strain T-22 and grown under salt stress. We determined the TRL of roots and analysed their polar metabolites using gas chromatography-mass spectrometry (GC-MS) and lipids using liquid chromatography-mass spectrometry (LC-MS) to provide new insights into the metabolism of inoculated and salt stressed roots compared to those grown in control conditions. This combined approach identified metabolic processes that are induced in barley during this interaction.

## Results

### Fungal association within roots confirmed using microscopy

Cultivar Gairdner roots were inoculated with *Trichoderma* strain T-22 and ink-vinegar staining was used to confirm the association of the fungus in colonized roots. Fungal spores were surrounding roots from day 2 until day 4, and on day 7 individual hyphae in heavily and partially colonized roots were clearly visible (Figure 1). For this reason, roots maintained in spore suspension for seven days were used for all further analyses.

**Figure 1.**
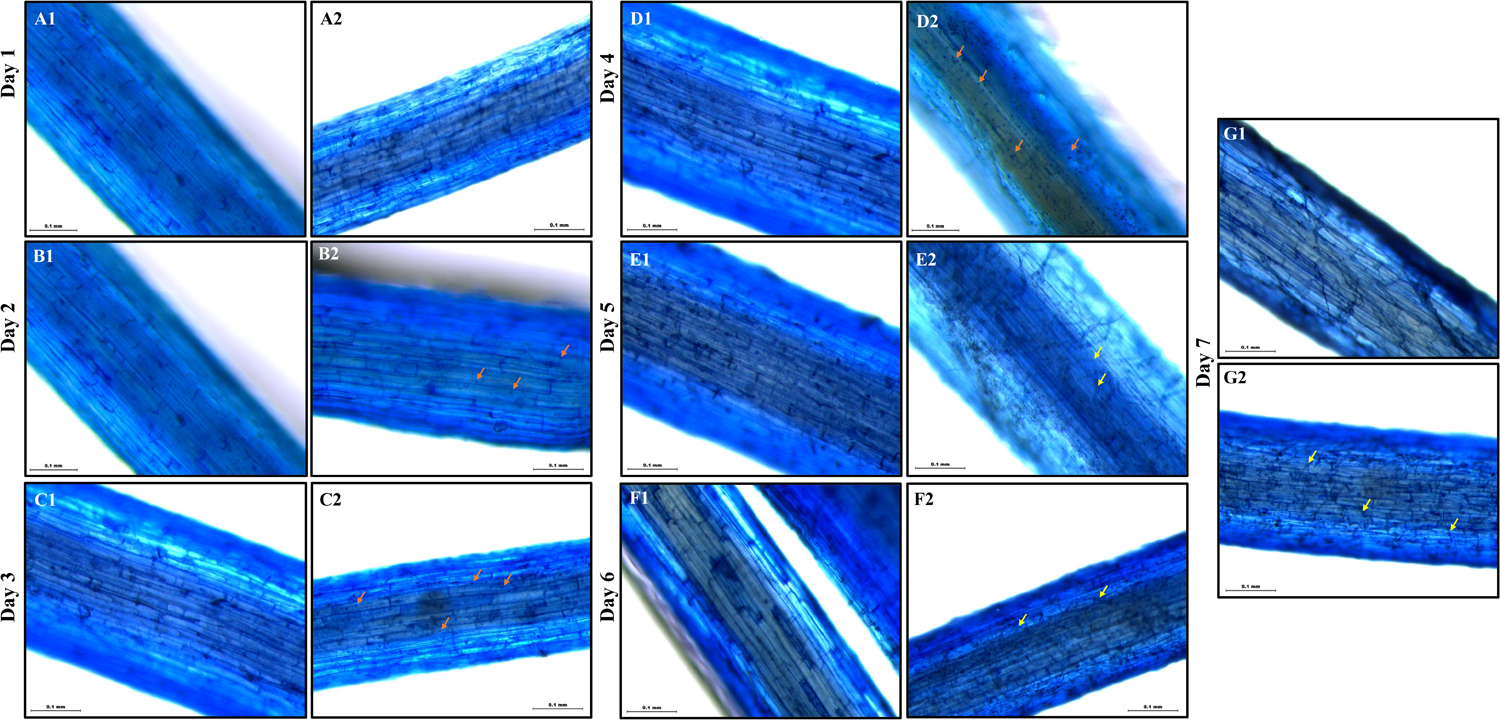
Light microscopy images of barley genotype cv. Gairdner over seven days. A1-G1, control barley roots; A2-G2, roots inoculated with *T. harzianum* T-22 (1 × 10^8^ CFU g^−1^ of seedling). Orange arrows in B2, C2 and D2 indicate fungal spores (dark blue spots); yellow arrows in E2, F2, and G2 indicate fungal hyphae (dark blue filaments). Scale = 0.1 mm. n = 3.

### Plant growth and root development

Two barley cultivars, Vlamingh and Gairdner, were selected based on their known differences in germination phenology and salinity tolerance. Vlamingh germinates well in high salt conditions while Gairdner is less tolerant (Gupta et al., 2019). Roots of uninoculated plants or those inoculated with *Trichoderma* strain T-22 (F) were treated with 0 (C) or 200 mM NaCl (S) and average TRL of each plant analysed using WinRHIZO root-scanning (Supplemental Figure S1).

Salt treatment of uninoculated plants (S) reduced TRL in both genotypes, but the reduction was more marked for Gairdner (Figure 2). In inoculated plants grown without salt (CF), TRL increased by 6.04% (*P* < 0.05) in Vlamingh and 9.87% (*P* < 0.01) in Gairdner compared to uninoculated roots of the same cultivar (C). In the inoculated salt treatment (SF), TRL increased by 14.85% (*P* < 0.05) in Vlamingh and 35.07% (*P* < 0.05) in Gairdner compared to treatment S.

**Figure 2.**
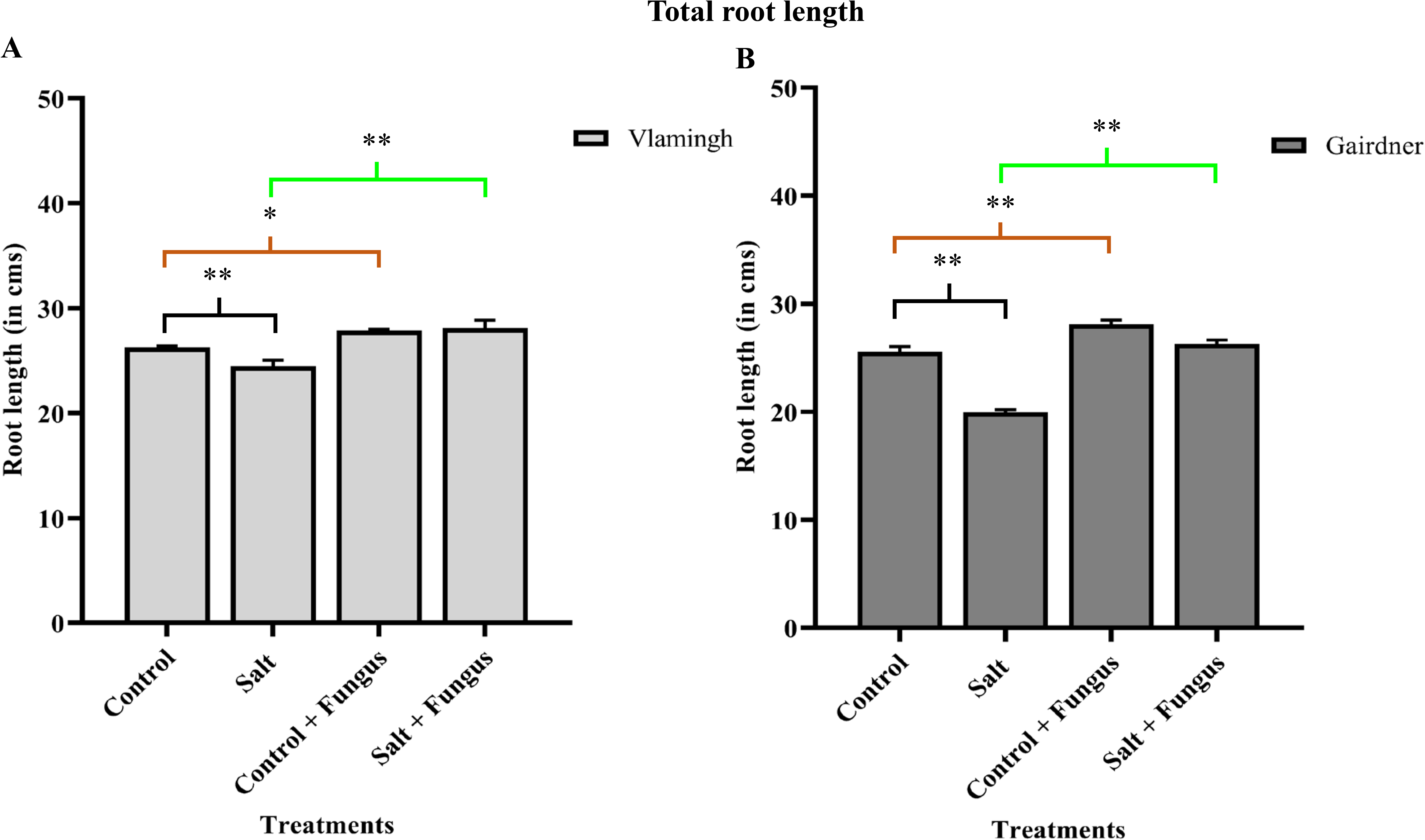
Total root length (TRL) of barley roots of two genotypes, (A) cv. Vlamingh and (B) cv. Gairdner, under four treatment conditions. X axis represent treatments and Y axis represent lengths in cm. n = 3. Coloured bars above graphs represent different comparisons: black indicates comparisons between salt and control: brown indicates comparisons between control + fungus and control; and green indicates comparisons between salt + fungus and salt. Asterisks denote significance for *P* < 0.05 (*) and *P* < 0.01 (**) as determined by Student’s *t*-test.

The decrease in TRL was 6.83% (*P* < 0.01) in Vlamingh and 23.73% (*P* < 0.01) in Gairdner when treatment S was compared to the same cultivar grown without salt (C). However, for inoculated plants, TRL of Vlamingh increased 0.91% (*P* < 0.01), while that of Gairdner decreased by only 6.24% (*P* < 0.01) after salt treatment (Figure 2).

### The effect of inoculation with *T. harzianum* and salt stress to polar metabolites in roots

Of the 82 metabolites detected in roots of both barley genotypes, 68 were identified using GC-MS (Supplemental Table S1). Of the known metabolites, 21 were amino acids (including amines), 20 were organic acids, 20 were sugars (including sugar derivatives) and five were fatty acids. The average normalized responses of all metabolites in all treated barley roots can be found in Supplemental Table S2.

A principal component analysis (PCA) of all metabolites showed a well-defined separation among samples from the different treatments (C, CF, S and SF) for both genotypes (Supplemental Figure S2a and S2b). The first and second principal components accounted for 53.5% and 50.3% of the variation in Vlamingh and Gairdner, respectively.

A heatmap representing the changes in metabolite levels in different treatments for both genotypes (Supplemental Figure 2c and 2d) shows a strong separation between the treatments for both genotypes. In Vlamingh, two major clusters were identified, with C, CF and S clustering while SF was separate. Further clustering was identified forming a subclass including C and CF while separating S. This implies fungal inoculation under salt stress had the strongest effect on the root metabolite profiles compared to all other treatments. In Gairdner, two major clusters were identified, the first including C and CF and the second including S and SF. The metabolites with the greatest variability amongst treatments were sugars.

The fold change of metabolites between the different treatments in both genotypes was analysed (Figure 3). In Vlamingh S, trehalose (+10.50-fold), arabitol (+5.30-fold), xylitol (+5.60-fold) and proline (+12.39-fold) showed the greatest increases compared to Vlamingh C. In Vlamingh CF, arabitol had the greatest increase, 3.04-fold, compared to Vlamingh C. In Vlamingh SF, sucrose (+3.13-fold), inositol (+6.60-fold), xylitol (+39.50-fold), ribitol (+15.60-fold), arabitol (+40.70-fold), rhamnose (+43.12-fold), ribose (+2.43-fold) and pipecolic acid (+2.20-fold) were strongly increased compared to S.

**Figure 3.**
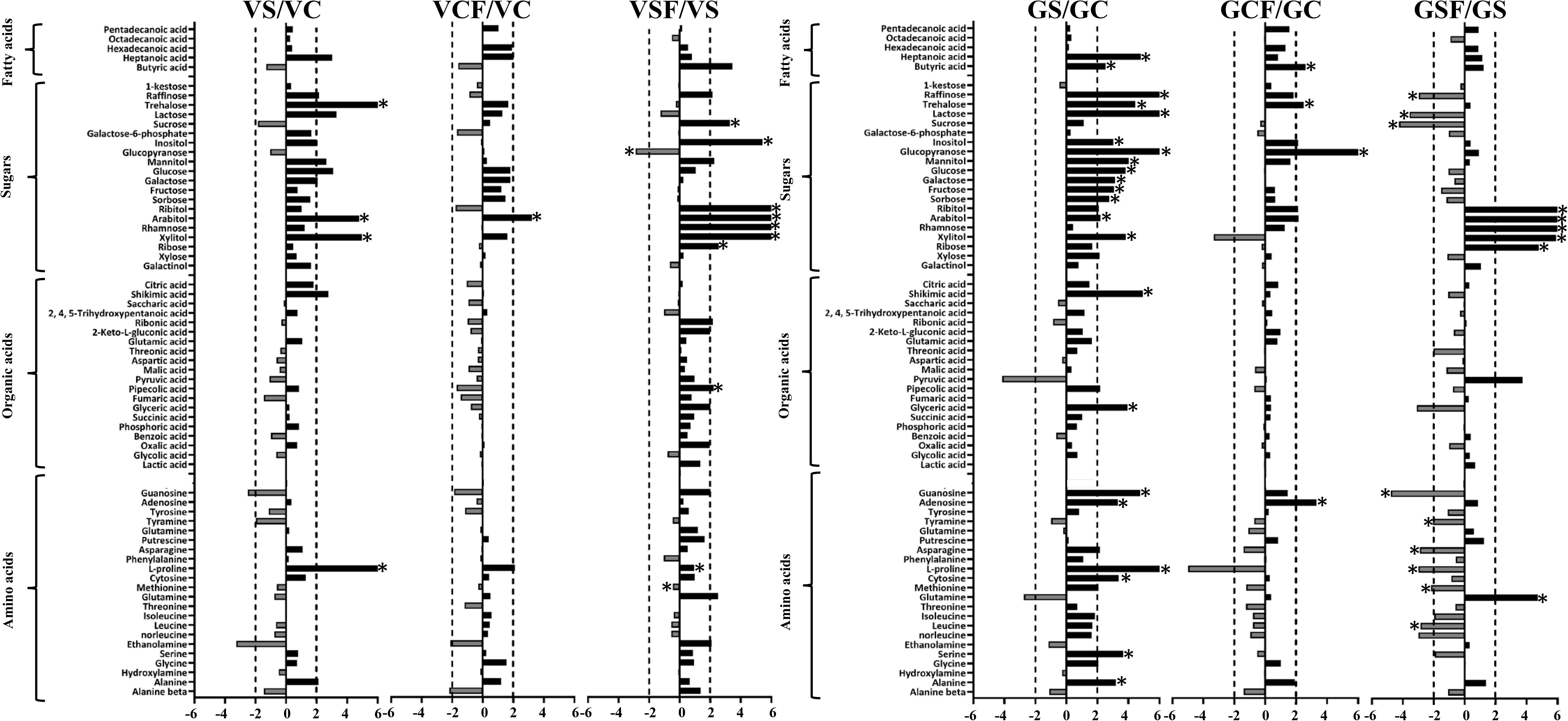
Logarithmic ratios of representative sugars, organic acids and amino acids in roots of barley cv. Vlamingh (V) and cv. Gairdner (G) of salt grown compared to control grown (VS/VC and GS/GC); control with fungus compared to control grown (VCF/VC and GCF/GC); and salt with fungus compared to salt grown (VSF/VS and GSF/GS). Values that are significantly higher (P < 0.05, FDR) are indicated by *. A threshold of ± 2-fold change is indicated by a dashed line.

In Gairdner S, 22 metabolites were increased with +2.0-fold or greater when compared to C and the highest were raffinose (+12.0-fold), trehalose (+4.67-fold), lactose (+8.60-fold), glucopyranose (+8.20-fold), shikimic acid (+5.52-fold), guanosine (+5.21-fold), and proline (+10.20-fold). In Gairdner CF, the metabolites that were increased most compared to C were trehalose (+2.40-fold), glucopyranose (+15.81-fold), adenosine (+3.14-fold) and butyric acid (+2.50-fold). In Gairdner SF, ribitol (+39.25-fold), arabitol (+91.20-fold), rhamnose (+73.80-fold), xylitol (+7.80-fold), ribose (+5.30-fold) and glutamine (+5.20-fold) were increased compared to S.

In Vlamingh SF, glucopyranose (−2.72-fold) decreased significantly compared to S. However, in Gairdner many more metabolites were decreased in SF compared to S, including raffinose (− 2.79-fold), lactose (−3.44-fold), sucrose (−4.35-fold), guanosine (−5.23-fold), tyramine (−2.02-fold), asparagine (−2.73-fold), proline (−2.82-fold), methionine (−2.12-fold) and leucine (−2.70-fold).

To elucidate the metabolic pathways involved and affected, identified metabolites were analysed using a pathway enrichment analysis. Supplemental Figure S3a shows an example of the metabolome view of all matched pathways arranged by *p* values on the Y axis and pathway impact values on the X-axis. Supplemental Figure S3b shows an example of a pathway view for alanine, aspartate and glutamate. The pathway given here is a simplified KEGG pathway and the matched metabolites are shown in red colour. Further detailed analysis identified biochemical pathways involved in the interaction of fungus with both genotypes. To achieve this, the number of identified metabolites were divided by the total number of metabolites in a pathway and a pathway coverage percentage was obtained. The top ten pathways affected in all treatments were the same for both genotypes (Figure 4)

**Figure 4.**
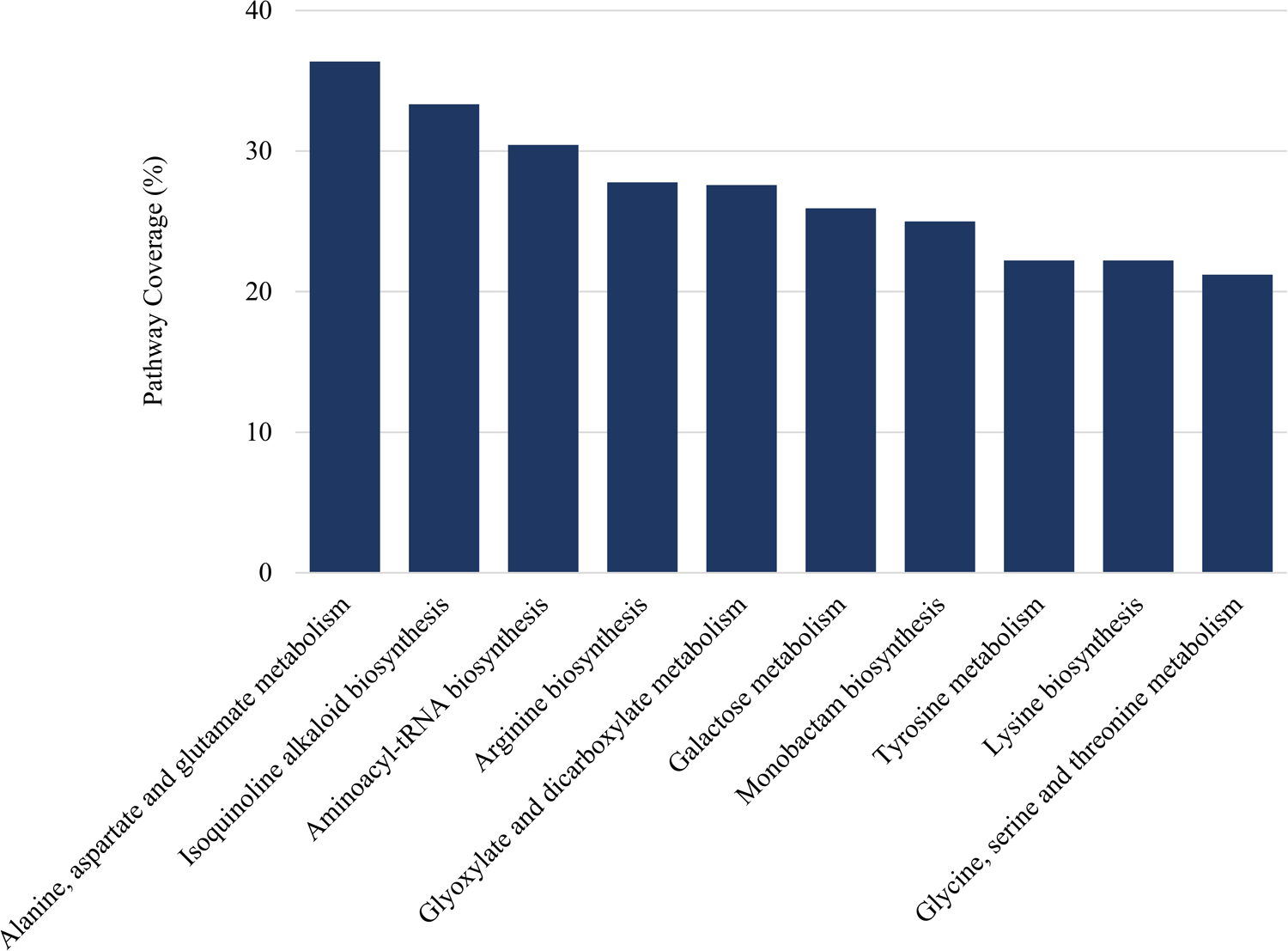
Pathway coverage graph for identified metabolites n roots of barley cv. Vlamingh and cv. Gairdner under four treatments. Top ten biochemical pathways that were affected are shown on X-axis and percentage of coverage is given on Y-axis.

### Global untargeted lipid profiling of barley roots

An untargeted LC-MS based study was performed to identify detailed changes in the lipidome of both barley genotypes in all treatments, C, CF, S and SF, and detected 3468 m/z_RT mass features (Table S3) which were subjected to multivariate analyses to identify discriminate differences.

PCA revealed clear separation between the treatments in both genotypes (Figure 5). The first and second components accounted for 72.3% of the variability between the 24 samples in Vlamingh and 63.6% in Gairdner. A hierarchical cluster analysis (HCA) heatmap (Supplemental Figure S4) showed a clear separation between treatments in both genotypes. The major clustering was between the control and salt treatments. Further sub-clusters were specific to genotypes with GSF clustered with GS and VSF clustered with VS. Under control conditions, VCF formed a separate cluster and VC, GC and GCF clustered together. Hence, clustering of the mass features was predominantly determined by the salt treatment followed by fungal treatment.

**Figure 5.**
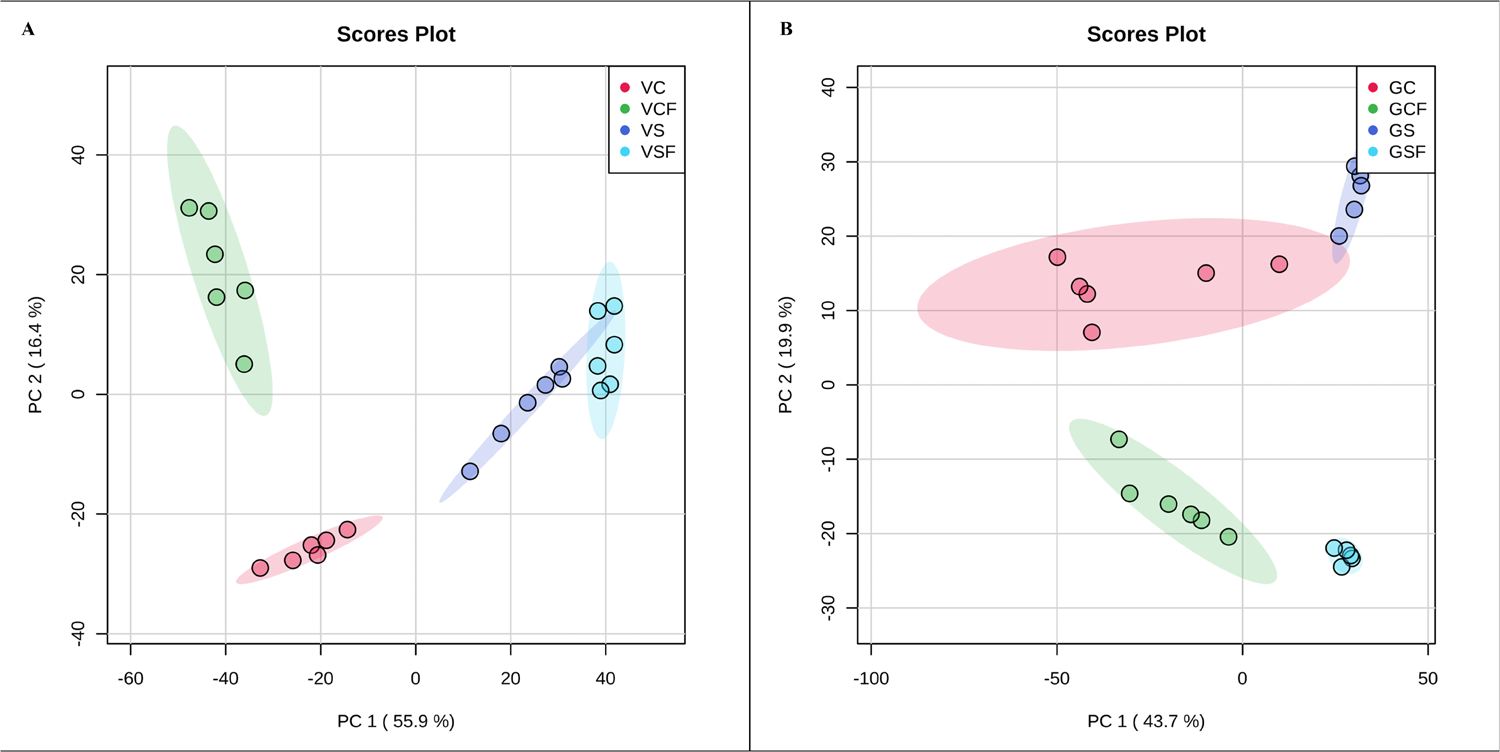
Principle component analysis (PCA) of the mass features (m/z_rt) extracted from lipid profiles of dissected roots of two barley genotypes grown under control and saline conditions with and without fungus. PC 1 versus PC 2 displays the separation between control and salt treatment for (A) *c*v. Vlamingh (V) and (B) cv. Gairdner (G). VC- control treated cv. Vlamingh; VCF- control and fungus treated *cv.* Vlamingh, VSF- salt and fungus treated *cv.* Vlamingh; GC- control treated *c*v. Gairdner; GCF- control and fungus treated cv. Gairdner, GSF- salt and fungus treated cv. Gairdner. n = 6 for all treatments. salt= 200 mM NaCl

There were much greater changes in the lipids in Vlamingh roots under saline conditions with 436 unique features showing significant changes compared to C, for Gairdner only 134 features were changed, while 107 lipids changed in both cultivars.

Fungus inoculated roots of Vlamingh under salt (VSF) exhibited only 58 significantly changed features compared to C while 176 changed in fungus inoculated roots of Gairdner after salt stress (GSF). 103 changed features were common to both genotypes (Supplemental Figure S5). Many more significantly different (*P* < 0.05) features were observed in Vlamingh and Gairdner (607 and 548 respectively) after short-term salt stress in plants not inoculated with fungus. This suggests that salt stress causes large changes in the lipid profile of roots but that inoculation with fungus greatly reduces the extent of these changes.

### Targeted MS/MS lipid analysis

MS-DIAL’s *in silico* LipidBlast database and an in-house plant lipid database were used to further identify lipids by the similarity calculation of retention time, precursor m/z, isotopic ratios, and MS/MS spectrum. False positives and true positives in the results for peak identifications were manually checked, curated and modified where needed. Of 3468 m/z_RT mass features, 522 m/z_RT mass features were annotated without MS2 spectra (Supplemental Table S4). 271 lipid molecular species were identified with MS2 spectra (Supplemental Table S5) which were categorised into different lipid classes (see Table 1). Lipid nomenclature follows the “Comprehensive Classification System for Lipids” given by the International Lipid Classification and Nomenclature Committee (Fahy et al., 2009). For example, the nomenclature PC (38:n) indicates a phosphatidylcholine (PC) species with a fatty acyl sum composition of 38 carbons containing all identified double bonds which are given as “n”. The information on the number of double bonds in each individual lipid species can be found in Supplemental Table S5.

**Table 1.**
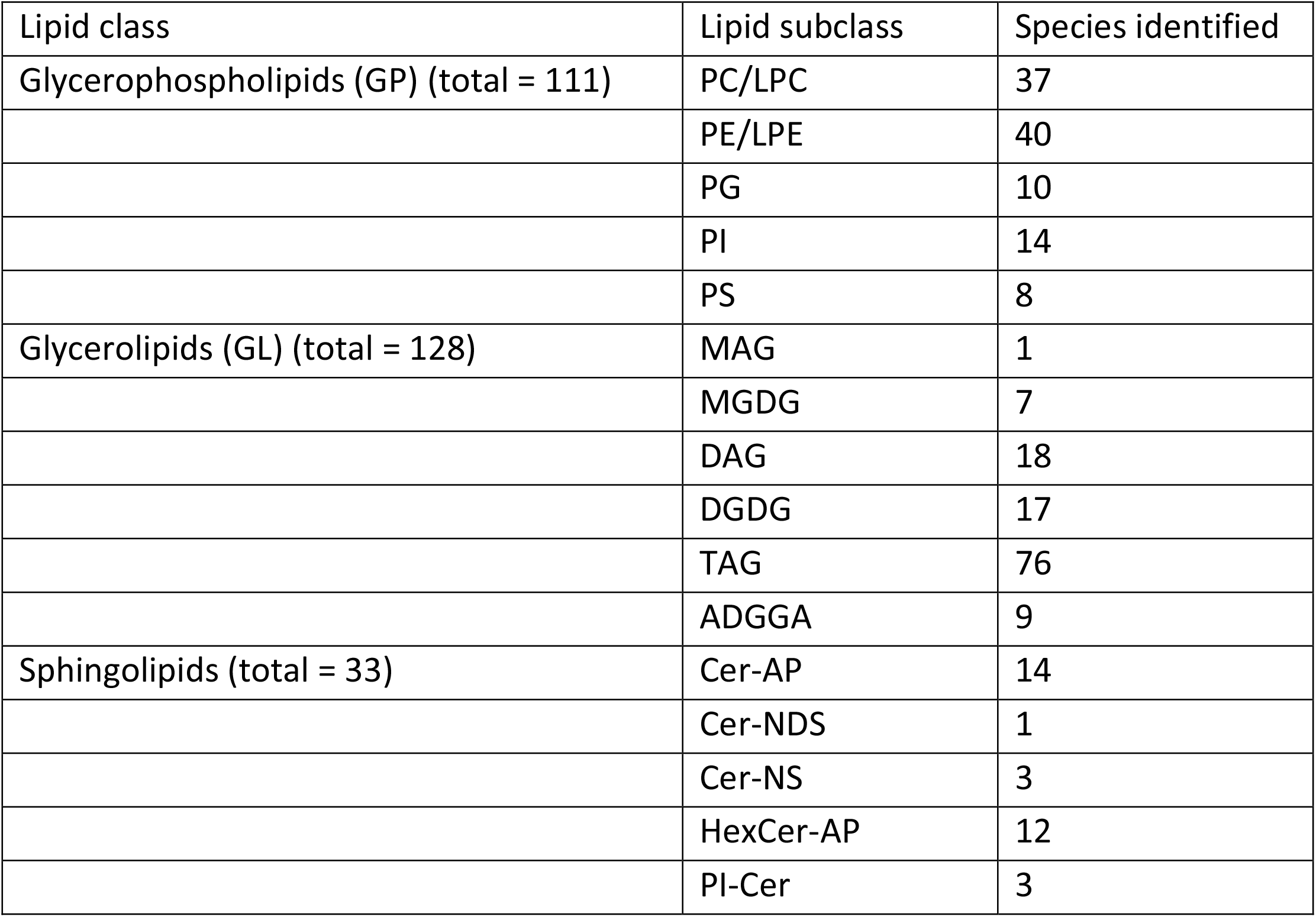
Summary of the number of identified lipid species from roots of two barley genotypes (cv. Vlamingh and cv. Gairdner) from four treatment groups- control C, control inoculated with fungus CF, salt (200 mM NaCl) S and salt inoculated with fungus SF. Analysis was performed using LC-TripleTOF-MS (positive ionization mode).

| Lipid class | Lipid subclass | Species identified |
| --- | --- | --- |
| Glycerophospholipids (GP) (total = 111) | PC/LPC | 37 |
|  | PE/LPE | 40 |
|  | PG | 10 |
|  | PI | 14 |
|  | PS | 8 |
| Glycerolipids (GL) (total = 128) | MAG | 1 |
|  | MGDG | 7 |
|  | DAG | 18 |
|  | DGDG | 17 |
|  | TAG | 76 |
|  | ADGGA | 9 |
| Sphingolipids (total = 33) | Cer-AP | 14 |
|  | Cer-NDS | 1 |
|  | Cer-NS | 3 |
|  | HexCer-AP | 12 |
|  | PI-Cer | 3 |

Pairwise comparisons between the four treatments in both genotypes were performed for Glycerophospholipids (GPs) and Glycerolipids (GLs) to determine the directionality of the changes (Table 2a and Table 2b).

**Table 2a.** Number of Glycerophospholipids as defined by Student’s t-test (*P* < 0.05, FDR) with a 2-fold change or greater, found for two barley genotypes in three treatments

| Lipid class | Treatment specific |  | Vlamingh | Gairdner |
| --- | --- | --- | --- | --- |
| Glycerophospholipids (GP) | Salt vs Control | Higher | 35 | 23 |
|  |  | Lower | 2 | 5 |
|  | Control+ fungus vs Control | Higher | 20 | 23 |
|  |  | Lower | 0 | 3 |
|  | Salt + fungus vs Salt | Higher | 0 | 35 |
|  |  | Lower | 11 | 7 |

**Table 2b.**
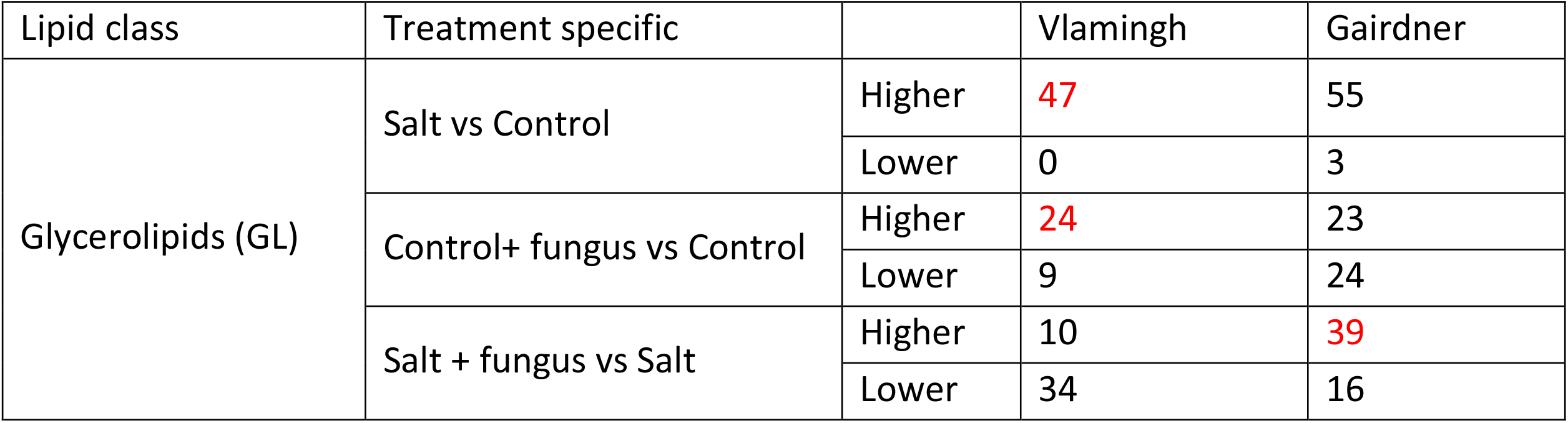
Number of Glycerolipids as defined by Student’s t-test (*P* < 0.05, FDR) with a 2-fold change or greater, found for two barley genotypes in three treatments

Vlamingh and Gairdner showed generally different responses with respect to GPs when treated with salt and/or inoculation. Salt treatment triggered a statistically significant increase in a much larger number of PC species in Vlamingh than in Gairdner. Of the 28 PCs, 19 increased in Vlamingh with significant fold changes greater than +2-fold, compared to changes in only four PC (42:n) in Gairdner (Figure 6a). PCs with a larger number of unsaturated bonds were particularly effected. PC (36:n) exhibited the strongest significant increase in Vlamingh with a +9.1-fold change after salt stress.

**Figure 6.**
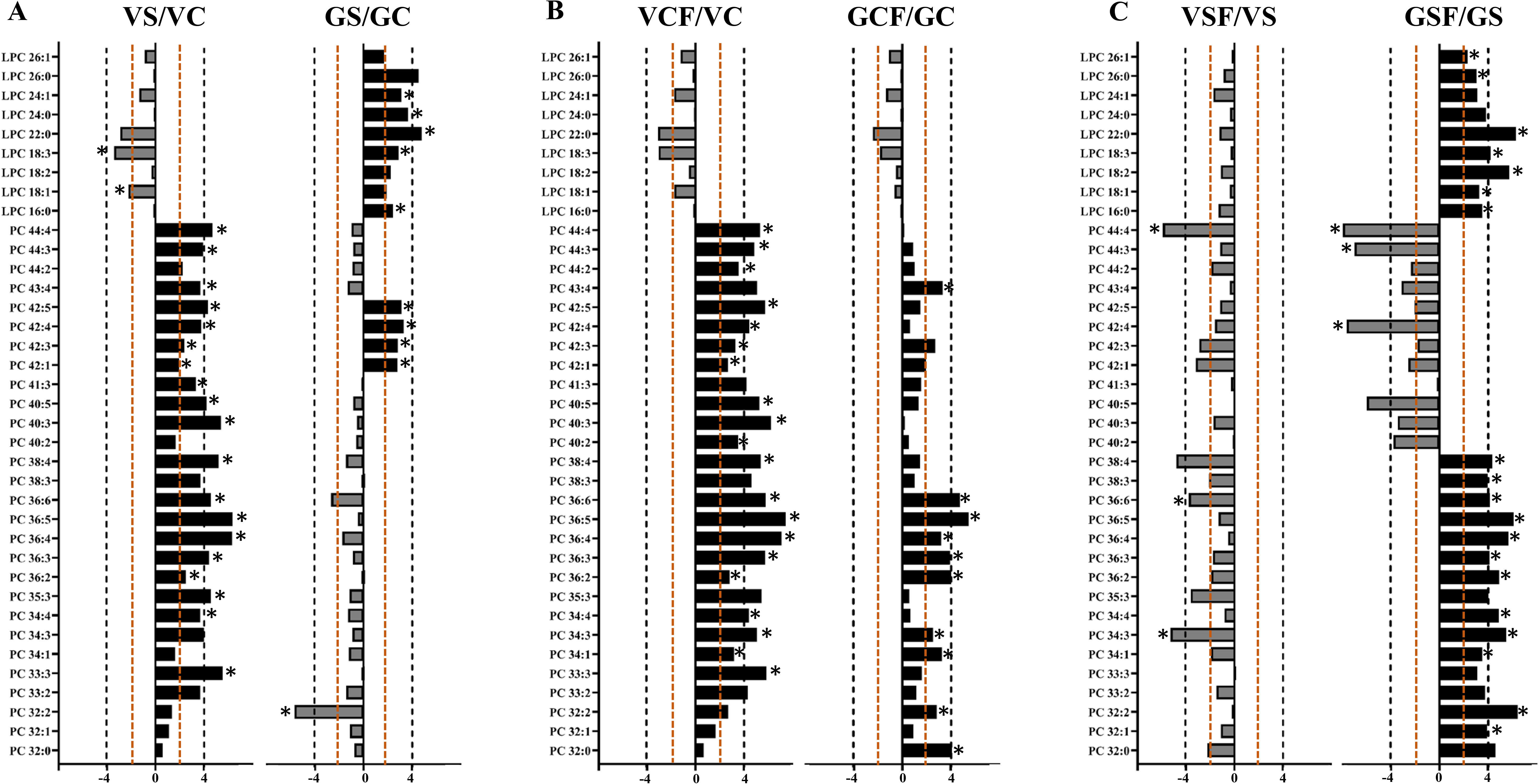
Logarithmic ratios of representative phosphatidylcholine (PC) and lysophosphatidylcholine (LPC) content in roots of barley. Cultivar Vlamingh and cv. Gairdner of salt grown (200 mM NaCl) compared to control grown; control with fungus grown compared to control grown; salt and fungus grown compared to salt grown. Values that are significantly higher (P < 0.05, FDR) are indicated by *. A threshold of ± 4-fold change is indicated by an orange dashed line and ±4-fold change is indicated by a black dashed line. X-axis represents the geometric progression with common ratio 4.

Inoculation with fungus had a similar effect to salt treatment in Vlamingh with 20 significant changes in PCs, 16 of those were the same as seen in the salt treatment, with PC (36:n) showing the largest fold change (Figure 6b). Fungal inoculation of Gairdner only caused significant increases in 10 PCs and these changes included an increase in the PC (36:n) lipids.

When the fungus inoculated Vlamingh roots were treated with salt there were few significant changes in PCs compared to the uninoculated plants illustrating the similar changes seen with both salt treatment and inoculation. The largest change was a decrease in PC (44:4). In contrast there was a large increase in PCs (32-38:n) in inoculated Gairdner compared to uninoculated after salt treatment. PC (44:4) was significantly decreased as in Vlamingh but PC (44:3) and (42:4) were also reduced (Figure 6c).

In Gairdner, salt treatment significantly increased LPCs and these were further increased in fungus inoculated roots in this treatment. This was in contrast to Vlamingh where LPCs did not change significantly in either salt treatment or after inoculation.

Salt treatment and/or inoculation with fungus caused few significant changes in LPEs in Vlamingh (Figure 7a, b, c). In Gairdner, fungal inoculation significantly reduced some LPEs (Figure 7b) but these were increased significantly when the inoculated plants were treated with salt (Figure 7c). Uninoculated plants had little changes in LPEs in response to salt (Figure 7a).

**Figure 7.**
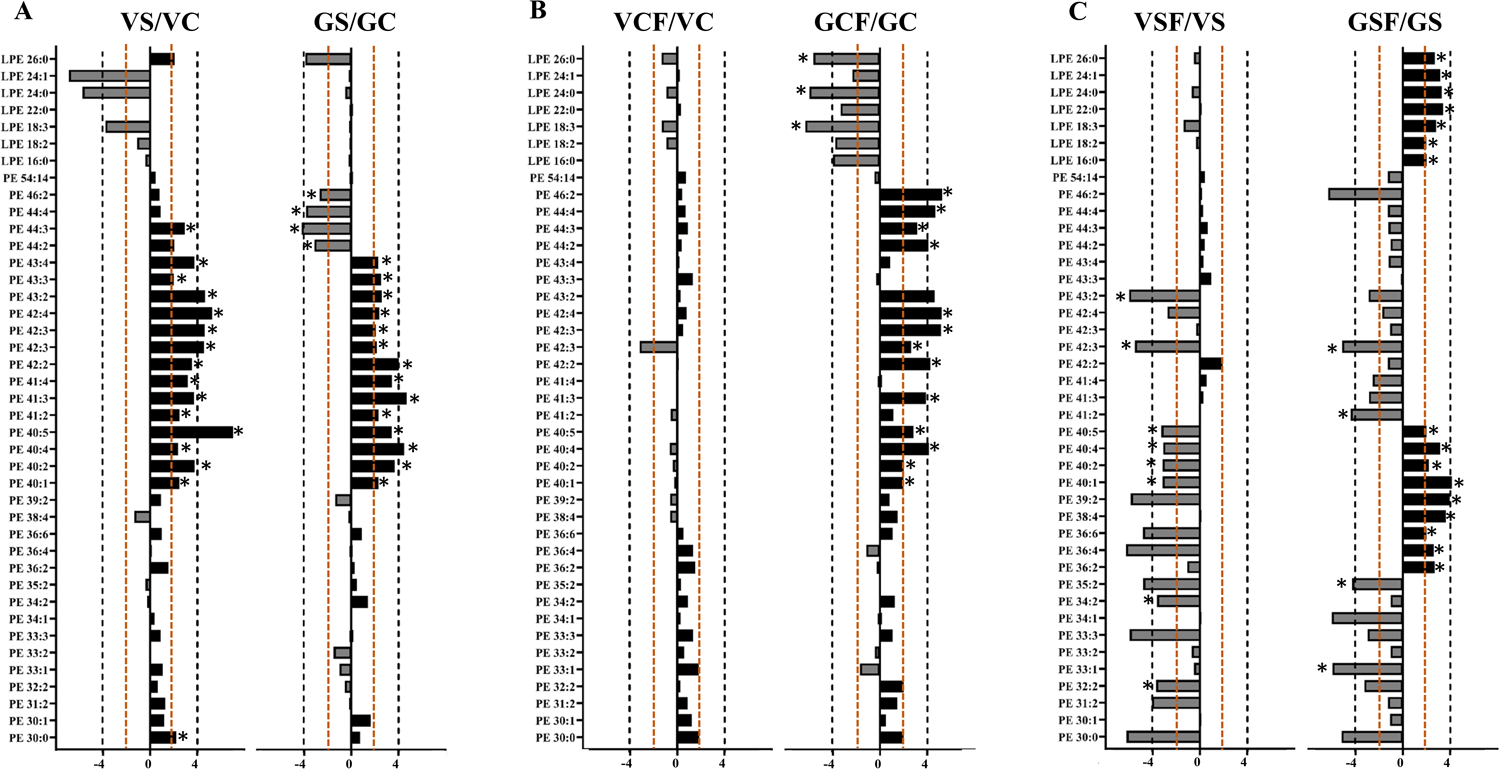
Logarithmic ratios of representative phosphatidylethanolamine (PE) and lysophosphatidylethanolamine (LPE) content in roots of barley. Cultivar Vlamingh and cv. Gairdner of salt grown (200 mM NaCl) compared to control grown; control with fungus grown compared to control grown; salt and fungus grown compared to salt grown. Values that are significantly higher (P < 0.05, FDR) are indicated by *. A threshold of ± 4-fold change is indicated by an orange dashed line and ±4-fold change is indicated by a black dashed line. X-axis represents the geometric progression with common ratio 4.

In both genotypes, salt stress significantly increased a large number of PE species. Sixteen PEs increased significantly after salt stress in Vlamingh and 14 in Gairdner. Compared to C, PE (40:n, 41:n, 42:n and 43:n) increased significantly under salt stress in both genotypes. PE (44:n) were significantly reduced in Gairdner but increased slightly in Vlamingh (Figure 7a, Supplemental Table S7). However, while 13 PEs (40:n, 41:3, 42:n, 44:n and 46:2) increased significantly in inoculated Gairdner roots, there were no significant changes in Vlamingh (VCF) (Figure 7b). Inoculation of Gairdner with fungus in control conditions (GCF) significantly increased PE (44:n) (Figure 7b, Supplemental Table S7) but these were significantly decreased in uninoculated plants after salt treatment (GS) (Figure 7a) and did not change in inoculated plants (GSF) (Figure 7c). In inoculated plants treated with salt (GSF), PE (40:n) further increased compared to uninoculated plants in the same conditions and PE (36:n, 38:4 and 39:2) also increased significantly (Figure 7c). For Vlamingh, eight PEs were significantly reduced in fungus inoculated roots subjected to salt treatment when compared to uninoculated plants (Figure 7c).

Salt had more pronounced effects on GL lipid species than inoculation with fungus. Of 128 glycerolipids, 75 increased by 2-fold or more in Vlamingh and 83 in Gairdner in response to salt (Figure 8a, 9a) whereas with fungal inoculation, only 24 of the 130 identified GLs increased in Vlamingh and 23 in Gairdner (Figure 8b, 9b). In Vlamingh, salt treatment of uninoculated roots (VS) increased five ADGGA lipids (Figure 8a, 9a) while in inoculated roots (VSF) six increased. However, many of these lipid species were significantly reduced in inoculated roots grown in normal conditions. In Gairdner, ADGGA showed a similar decrease after fungal inoculation (GCF) without salt treatment but increased after salt treatment (GFS).

**Figure 8.**
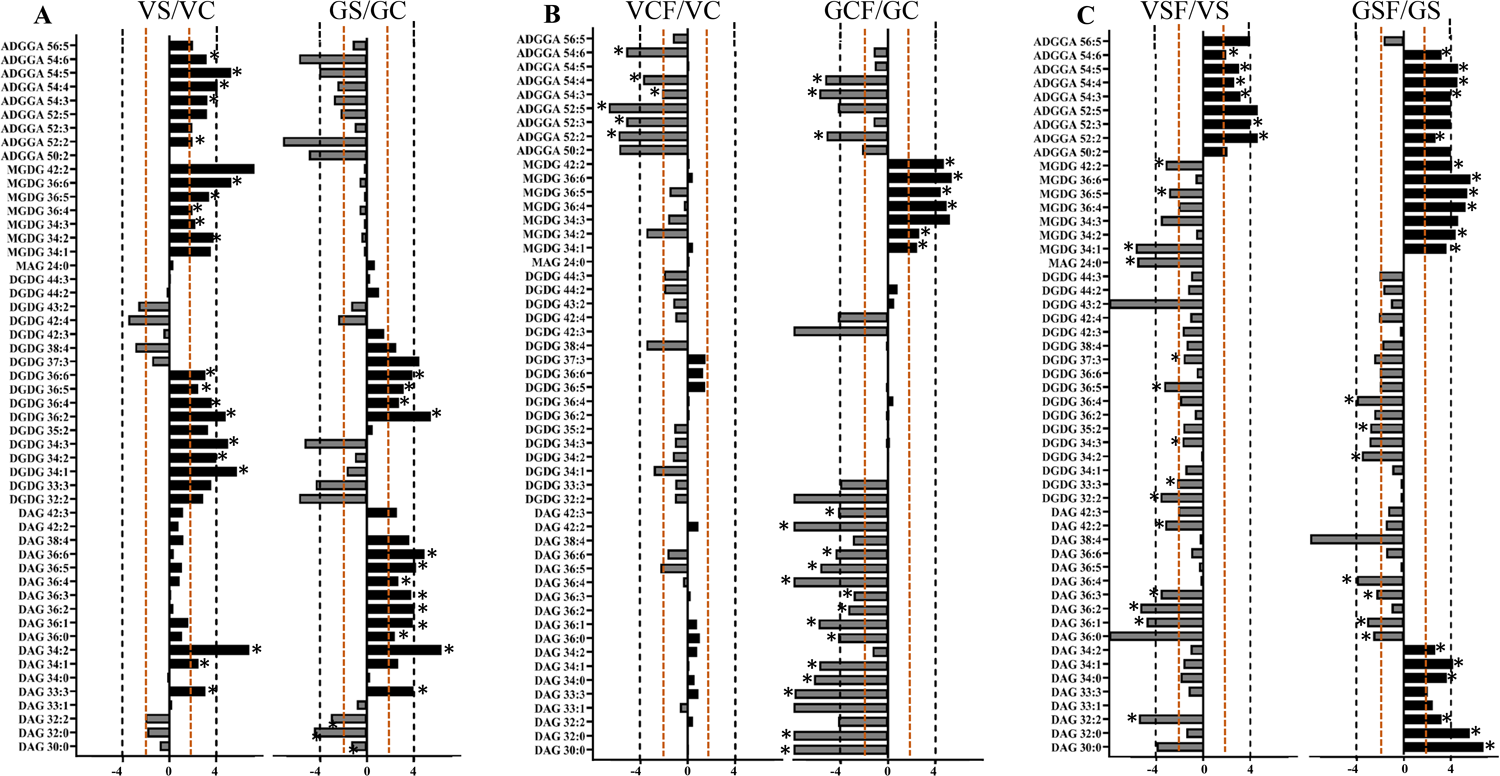
Logarithmic ratios of representative acyl diacylglyceryl glucuronide (ADGGA), monogalactosyl diacylglycerol (MGDG), digalactosyl diacylglycerol (DGDG), monoacylglycerol (MAG) and diacylglycerol (DAG) content in roots of barley. Cultivar Vlamingh and cv. Gairdner of salt grown (200 mM NaCl) compared to control grown; control with fungus grown compared to control grown; salt and fungus grown compared to salt grown. Values that are significantly higher (P < 0.05, FDR) are indicated by *. A threshold of ± 4-fold change is indicated by an orange dashed line and ±4-fold change is indicated by a black dashed line. X-axis represents the geometric progression with common ratio 4.

**Figure 9.**
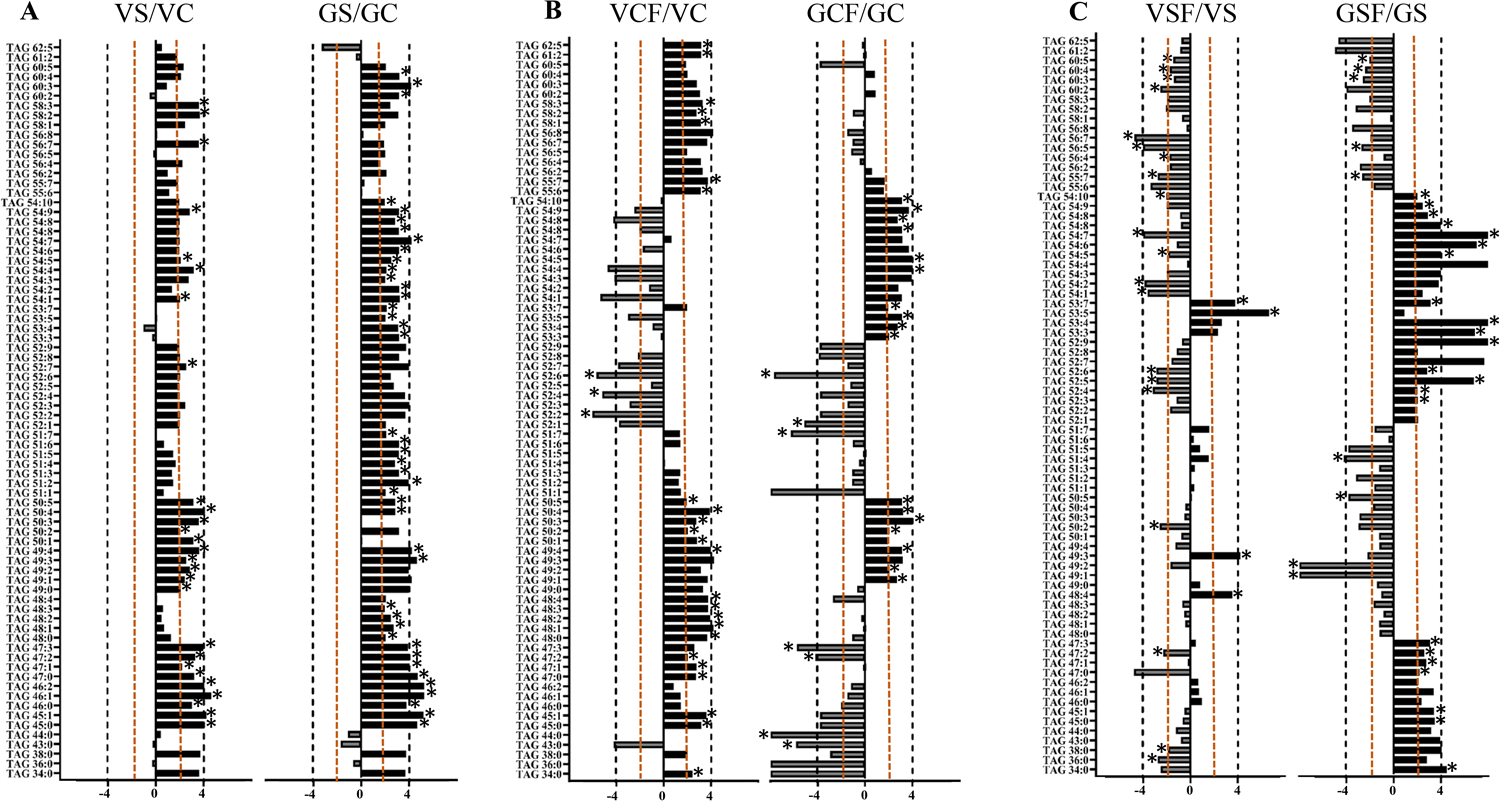
Logarithmic ratios of Triacylglycerol (TAG) content in roots of barley. Cultivar Vlamingh and cv. Gairdner of salt grown (200 mM NaCl) compared to control grown; control with fungus grown compared to control grown; salt and fungus grown compared to salt grown. Values that are significantly higher (P < 0.05, FDR) are indicated by *. A threshold of ± 4-fold change is indicated by an orange dashed line and ±4-fold change is indicated by a black dashed line. X-axis represents the geometric progression with common ratio 4.

In Vlamingh, but not Gairdner, MGDG increased in response to salt treatment (VS) (Figure 8a) but were reduced in roots of salt treated inoculated plants (VSF) (Figure 8c). In contrast, these lipids were not increased in salt treated Gairdner roots (GS) (Figure 8a) but increased after fungal inoculation (GCF, GSF) (Figure 8b, c).

Of the 17 DGDG species, seven showed significant increases in salt stressed roots of Vlamingh, while only four increased significantly in Gairdner. DGDG (36:n) increased significantly in both genotypes. Of seven identified MGDGs, five increased significantly in Vlamingh but not in Gairdner (Figure 8a, Supplemental Table S8). Fungal inoculation had little effect on DGDG species (VCF, VSF, GCF, GSF) (Figure 8b, c) although a small number decreased significantly in inoculated roots treated with salt (VSF, GSF) (Figure 8c).

In Vlamingh, only three DAGs were increased under salt while nine DAGs exhibited significant increases in Gairdner (Figure 8a, Supplemental Table S8). Fungal inoculation significantly decreased 14 DAGs in Gairdner roots but there were no significant changes in Vlamingh (Figure 8b). In salt treated roots of inoculated plants five DAGs were reduced significantly in Vlamingh. In Gairdner four DAGs were reduced significantly and six increased significantly (Figure 8c).

TAGs were the GL species most affected by salt stress with 27 TAGs exhibiting significant increases in Vlamingh roots (VS) and 42 in Gairdner (GS) (Figure 9a, Supplemental Table S9). Fungal inoculation affected 24 TAGs with unsaturated double bonds in Vlamingh with some increasing and others decreasing (Figure 9b). In Gairdner, 17 TAGs were significantly increased following inoculation and seven significantly decreased (Figure 9b, Supplemental Table S9). Plants inoculated with fungus in the salt treatment (GSF) had increased levels of 22 TAGs (54:n, 53:n, 52:n, 47:n, 54:n and 34:n) compared to control (GS) and reduced levels of nine (49:1, 49:2, 50:n, 51:n and 60:n species) (Figure 9c). In contrast, 19 TAGs were significantly reduced in Vlamingh inoculated plants treated with salt and only four were increased (Figure 9c).

## Discussion

In this study we have shown the beneficial effect of inoculation of barley roots with *T. harzianum* T-22 on TRL in both control and saline conditions. We confirmed that the effect of salinity was more severe in the sensitive genotype Gairdner than in the tolerant genotype Vlamingh. Inoculation with fungus in saline conditions improved the TRL of both Vlamingh (14.85%) and Gairdner (35.07%) compared to uninoculated plants, resulting in equivalent values to those of inoculated and uninoculated roots grown in normal conditions. These results demonstrate the ability of the fungus to mitigate the negative effects of NaCl, results that are consistent with the findings of Mastouri et al. (2012).

Analysis of metabolites and lipids that changed during inoculation with fungus and in response to salt treatment gave insights into the mechanisms by which inoculation with *T. harzianum* T-22 was able to help the plant respond to saline conditions. It showed that fungal inoculation during salt treatment had a more significant effect on metabolites, particularly sugars, while the changes in lipid composition were more an effect of salt treatment than the inoculation. Gairdner showed more obvious lipid remodelling than Vlamingh, but for both the fungus mostly affected neutral lipid species.

### Fungal inoculation promoted root growth

Total Root Length (TRL) was measured as a means of determining plant development. Salinity reduced root length of both genotypes. According to Acosta-Motos et al. (2017) and Shelden et al. (2016), this could be due to inhibitory impacts of salts, restricting cell division and cell expansion in the root apical meristem. The effect of salinity was more severe in the sensitive genotype Gairdner than in the tolerant genotype Vlamingh, confirming the greater salt tolerance potential of the latter. Fungal inoculation appeared to have a general growth promotion effect on roots (Contreras-Cornejo et al., 2009) but this was more significant when the plants were grown in saline conditions.

### Fungus changes osmolytes in roots as a protective adaptation to salt stress

In this study, a global analysis of the GC-MS dataset using PCA and HCA indicated substantial osmolyte differences between the two inoculated and uninoculated barley genotypes under control and saline conditions. More sugars changed significantly in Gairdner than in Vlamingh after salt stress suggesting the higher need for balancing the osmotic pressure within the cell and stabilizing the cell membrane in the more sensitive genotype. Sugars provide carbon and energy for normal functioning of cellular metabolism and regulate growth and development of plants. Sugars and sugar alcohols (polyols) have been widely accepted as osmoprotectants (Singh et al., 2015). Similar results were shown by (Cao et al., 2017) where varying changes in sugar metabolism after salinity were detected for several barley genotypes. Compared to Vlamingh, more significant changes were observed in polyols in roots of Gairdner after salt stress. It has been suggested that sugar alcohols such as mannitol, xylitol and ribitol enhance tolerance of plants against drought and salinity (Williamson et al., 2002). Increased synthesis of these osmoprotectants helps plants to quench the most toxic hydroxyl radical (OH) generated via the Fenton reaction (Gill and Tuteja, 2010). Therefore, it can be suggested that in Gairdner there was an increased requirement for water adjustment in the cytoplasm, Na^+^ sequestration into the vacuole or apoplast, and protection of cellular structures than in Vlamingh.

Under control conditions, sugars were not increased in either genotype inoculated with *T. harzianum*. This may be the result of the activation of one of the major starch-degrading enzymes, glucan-water dikinase, by the fungus in colonized roots as reported by Sherameti et al. (2005). Similar results were observed by Ghabooli (2014) in fungus inoculated plants. However, fungal inoculation increased levels of sugars in both genotypes under salt which may help roots to protect against the severe effects of salinity and thus increase the growth rate of inoculated roots.

The salt sensitive genotype, Gairdner, showed larger significant increases in amino acid levels compared to Vlamingh under salt stress (uninoculated). However, the fungus inoculated roots showed significantly decreased amino acid levels in Gairdner in saline conditions. Here we can speculate that the fungus may enhance the capacity of Gairdner to withstand salt stress through coordinating with amino acids exerting a protective mechanism or a significant reduction in amino acid levels could indicate their contribution to growth in Gairdner (inoculated) under saline conditions. This is also in alignment with past research where Li et al. (2017) found that *Aspergillus aculeatus* inoculated roots of ryegrass showed decreased amino acid concentrations following salt stress.

### Alterations in the lipid profiles in roots of Vlamingh and Gairdner under various treatments

Untargeted lipidome analyses were performed on roots of both genotypes to elucidate the role of lipid species in adaptive mechanisms following short-term salt stress. The untargeted data strongly suggests that salt treatment has a much bigger impact on the root lipidome than the effect of fungus in both genotypes. However, there were some changes related to fungal inoculation that may be linked to better performance of barley roots in saline conditions.

A number of lipid classes show patterns of changes that may be associated with tolerance to salt. These include PC, ADGGA and MGDG. Vlamingh, which is tolerant with only a small change in TRL in response to salt, increases of these lipids may be part of its response, while the sensitive Gairdner, which shows a much larger reduction in TRL, does not. Although these lipid species show differing responses to inoculation in the two varieties, in all cases the levels of these lipids are significantly increased in inoculated Gairdner in response to salt treatment and this is accompanied by only a small reduction in TRL (compared to inoculated plants).

For example, fungal inoculation of Vlamingh significantly increases the same PCs that are increased in response to salt and these are mostly maintained at a similar level after salt treatment. TRL does not change significantly after salt treatment of inoculated plants, and in fact it is slightly higher than uninoculated plants grown in control condition. In contrast, Gairdner only increases a small number of PCs in response to salt treatment and its TRL shows a large reduction. After fungal inoculation, it increases many more PCs and, in these roots, the TRL is significantly higher than in roots grown in control conditions. Salt treatment of inoculated Gairdner increases PC 32-36:n and PC 38:n. The changes in PC levels may relate more to improved growth as PCs stay at the same level in inoculated roots grown in normal conditions and in salt.

For Vlamingh, the pattern of increase in ADGGA species is different to that of PCs. Although they are increased in uninoculated and inoculated roots in response to salt, they are not increased in response to inoculation. This suggests that they are part of the tolerance responses. Gairdner does not increase ADGGAs in response to salt unless it is inoculated suggesting that the fungus changes the ability of Gairdner to deal with salt stress by upregulating the synthesis of these lipid species. To the best of our knowledge, we found that the role of ADGGA is mostly reported against phosphorus depletion as also observed by Okazaki et al. (2013) in Arabidopsis. However, in our study, there is a clear association of ADGGA with salt tolerance. Therefore, this needs to be further studied.

In uninoculated plants, MGDG lipid species also increase in Vlamingh in response to salt treatment but after inoculation this response is not so marked (there are lower levels of some MDGD in inoculated salt treated roots compared to control salt treated roots). However, inoculation increases MGDG species in Gairdner and the same lipids are increased after salt treatment in inoculated roots but not uninoculated suggesting they may be part of the response that allows the inoculated Gairdner to tolerate salt better than uninoculated roots.

Diacylglycerol is known to be an intermediate in the synthesis of membrane lipids and is involved in phospholipid signalling in plant cells (Dong et al., 2012). Plants can utilize at least two metabolic pathways to produce different molecular species of the immediate precursor to TAG, diacylglycerol (DAG): (1) *de novo* DAG synthesis (termed as DAG1 here), and (2) conversion of the membrane lipid phosphatidylcholine (PC) to DAG (termed as DAG2 here). Here, since there was no significant increase or change in PC species in Gairdner after salt stress, it can be postulated that DAG and subsequent TAG production was through utilization of the products of FA synthesis exported from the plastid directly for *de novo* DAG/TAG synthesis through the Kennedy pathway (Weiss and Kennedy, 1956) (Figure 10a). Conversely, in Vlamingh, TAG synthesis was presumably through PC-derived DAG/TAG synthesis (Figure 10b). The pathway of PC-derived DAG2 synthesis starts with the production of PC lipids from *de novo* DAG1. PCs are synthesized through CDP-choline:diacylglycerol cholinephosphotransferase (CPT) (Li-Beisson et al., 2013) leading to a net production of PCs from DAGs. Further, phosphatidylcholine:diacylglycerol cholinephosphotransferase (PDCT) (Lu et al., 2009) activity transfers the phosphocholine head group from PCs to DAGs generating new molecular species of DAGs and PCs, which does not lead to net accumulation of PCs or DAGs. This supports our findings where only three DAGs increased after salt stress in Vlamingh with nine DAGs in Gairdner. Once PC lipids are formed from *de novo* DAG1 by either CPT or PDCT, the FAs esterified to PCs are available for FA modification and acyl editing (Bates et al., 2009; Li-Beisson et al., 2013) to generate new molecular PCs. The DAG substrate for TAG synthesis is subsequently derived from PCs by removal of the phosphocholine headgroup (Bates, 2012). This eventually results in TAG production.

**Figure 10.**
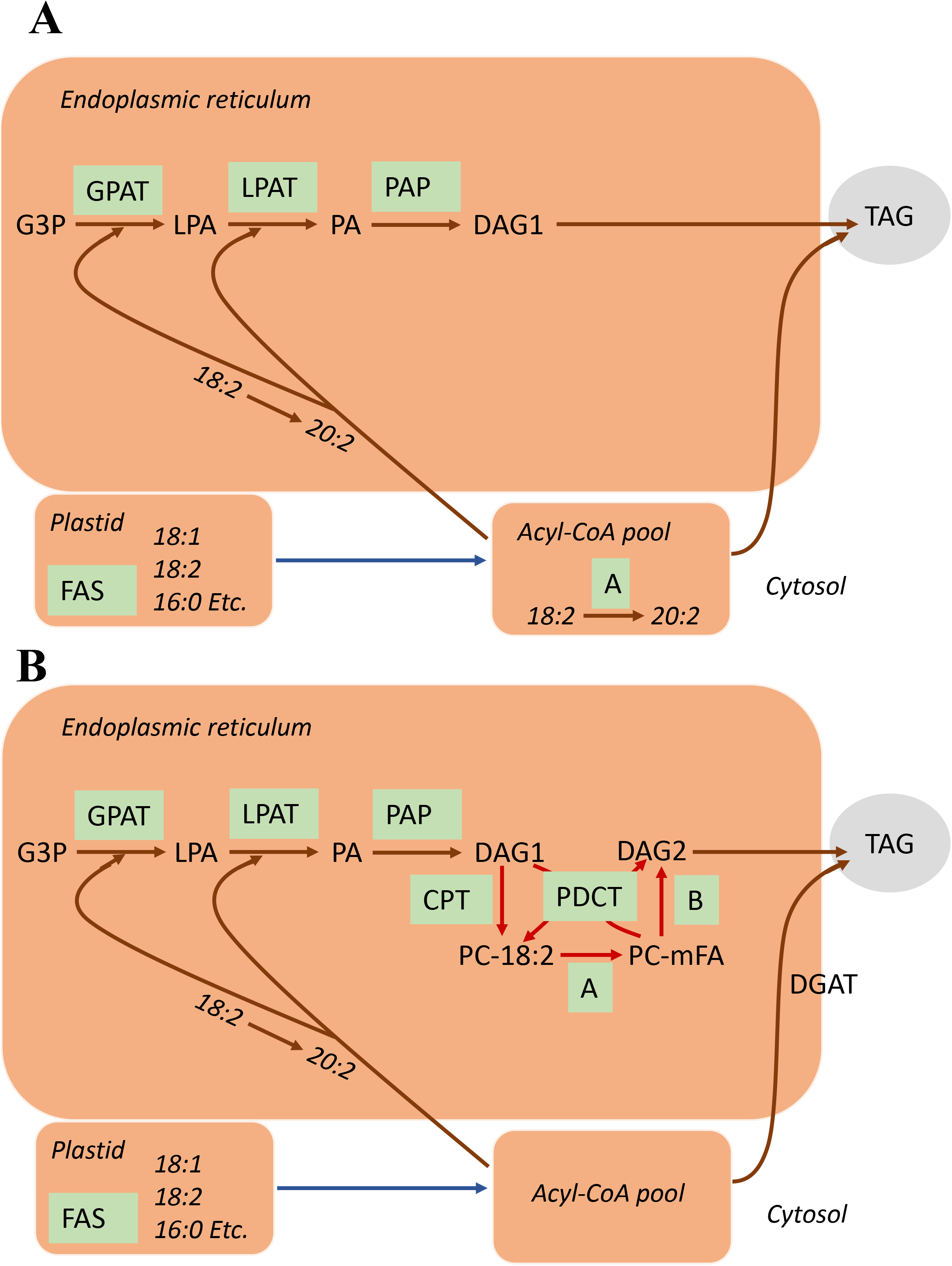
*De novo* TAG synthesis using the Kennedy pathway (A) and PC-derived TAG synthesis. Brown arrows indicate reactions involved in *de novo* DAG1 synthesis (B). A: Brown arrows indicate reactions involved in *de novo* DAG1 synthesis. Enzymatic reactions are embedded in green boxes: A- FA elongation, DGAT- acyl-CoA:DAG acyltransferase, FAS- fatty acid synthesis, LPAT- acyl-CoA:LPA acyltransferase, GPAT- acyl acyl-CoA:G3P acyltransferase, PAP- PA phosphatase. Substrates are DAG- diacylglycerol, G3P- glycerol-3-phosphate, LPA- lyso- phosphatidic acids, PA- phospatidic acid. B: Red arrows indicate reactions involved in PC- derived DAG2 synthesis. Enzymatic reactions are embedded in green boxes: A- modification of FA esterified to PC, B- reversible CPT or phospholipase C or phospholipase D/PAP DAG production. CPT- CDP-choline:DAG cholinephosphotransferase, DGAT- acyl-CoA:DAG acyltransferase, FAS- fatty acid synthesis, LPAT- acyl-CoA:LPA acyltransferase, GPAT- acyl acyl- CoA:G3P acyltransferase, PDCT- PC:DAG cholinephosphotransferase, PAP- PA phosphatase. Substrates are DAG- diacylglycerol, G3P- glycerol-3-phosphate, LPA- lyso-phosphatidic acids, PA- phosphatidic acid.

Under control conditions, *T. harzianum* inoculation did not change the DAG profile in roots of Vlamingh. But a significant increase in TAGs in inoculated roots of Vlamingh suggests the ability of the fungus to synthesize fatty acids *de novo* and incorporate them into TAGs through the Kennedy pathway as shown in Figure 10a (Weiss and Kennedy, 1956). A significant reduction in DAGs in inoculated roots of Gairdner suggests a role of DAGs in the synthesis of TAGs which were strongly increased after inoculation in Gairdner.

An interesting pattern with reduction in higher numbers of neutral lipids was observed in inoculated roots of Vlamingh after salt stress. Downregulation of TAGs was proposed to be an adaptive feature of salt-tolerant plants by (Barkla et al., 2018) providing precursor lipid molecules such as DAG and FA for phospholipid synthesis. Therefore, it can be suggested that fungus imparts tolerance to Vlamingh by reducing TAGs under saline conditions.

Under saline conditions, TAGs accumulated in inoculated roots of Gairdner. Here, higher accumulation of TAGs may reflect lipid remodelling, i.e. channelling of fatty acids from constitutive membrane lipids to TAGs. Further, as TAGs predominantly contain polyunsaturated fatty acids, it is likely that the fungus helps in the release of polyunsaturated fatty acids from structural lipids to be used for transient assembly of TAGs during membrane remodelling. Similar results were observed by (Mueller et al., 2015) where salt stress triggered accumulation of TAGs in Arabidopsis seedlings.

There appears to be a pattern that Vlamingh, which is tolerant to salt and with little reduction in TRL, is innately able to modify its lipids to respond but Gairdner requires the fungus to be able to initiate this response. For this reason, inoculation of Gairdner has a much more significant effect combating negative effects salt stress. Inoculation of both varieties increases the TRL of roots and this may allow them to better tolerate salt treatment. More research is required to clarify the links we have identified between improved growth, changes in lipid and metabolite profile and salt tolerance.

### Conclusions

This study investigated the role of *Trichoderma harzianum* T-22 on two barley genotypes, Vlamingh and Gairdner, that differ in their germination phenology and salinity tolerance. Total root length was measured after the final harvest. We showed that salt treatment has an effect on TRL but the fungus allows the roots to maintain their growth better, confirming the positive role of the fungus to help roots cope with the adverse effects of salt stress. We investigated the metabolite and lipid changes in roots of both genotypes, using untargeted GC-MS and LC-MS approaches allowing unique insights into the metabolome and lipidome of roots influenced by the inoculation of endophytic fungi identifying metabolic processes that are induced during this interaction. Based on our results, the fungus modifies the metabolome and lipidome of each genotype in a specific manner, improving their ability to cope with salt stress. Fungus-plant interactions and their effect on plant growth and metabolism are important questions which our study aims to address. The fungus was unable to promote equivalent levels of tolerance in both genotypes although it provided significant improvement in growth of the sensitive genotype Gairdner.

## Materials and Methods

### Plant Materials and growth conditions

Two barley genotypes (Vlamingh and Gairdner) were selected based on their known differences in germination phenology and salinity tolerance (Gupta et al., 2019). All seeds were sourced from the University of Adelaide, Australia.

### Fungal isolates and growth conditions

Freeze dried form of endophyte *Trichoderma harzianum Rifai* strain T-22 (ATCC 20847) was sourced from The American Type Culture Collection (ATCC, Washington, DC). The pellet obtained in the ampoule was suspended with sterile distilled water and kept at room temperature (25°C) undisturbed overnight. On the following day, the suspension was mixed well and inoculated on Potato Dextrose Agar (39 g/L PDA powder in de-ionized water) in solid medium on a petri plate. The plates were incubated at 25°C in dark for 5 days followed by light for 3 days. Conidia was seen growing after 8 days of incubation. The plates were stored at room temperature prior to setting up the symbiosis interaction.

### Material and chemicals

Analytical or mass spectrometric grades solvents and reagents were purchased from Merck Millipore (Bayswater, VIC, Australia). 18.2 Ω deionized water was obtained from Synergy UV Millipore System (Millipore, USA). Lysing Matrix Tubes with 0.5 g Lysing Matrix D (1.4 mm ceramic spheres) were purchased from MP Biomedicals (Seven Hills, NSW, Australia). PDA was purchased from Sigma-Aldrich.

### Plant growth conditions and inoculation with *T. harzianum*

A flow-chart showing the experimental setup and a detailed description of for plant growth conditions and inoculation with fungus are provided in Supplemental Figure S6 and Supplemental Materials and Methods. In brief, 24 seeds from each variety were sterilized and washed before imbibition (overnight ~16 h). Then, seeds were transferred to agar plates containing Whatman paper soaked in 5ml 18.2 Ω deionized water. Plates were sealed, wrapped in aluminium foil and kept in a growth chamber maintained at 17°C constant temperature with no light. Plates were unwrapped after 48 hours of germination period and transferred to a growth chamber maintained at 17°C for 16 hours light and 10°C for 8 hours dark cycles for five days.

After seven days, 12 seedlings from each variety were inoculated fungus and 12 control seedlings from each variety were treated with an equal amount of sterile water. Seedlings were kept at 17°C for 16 hours light and 10°C for 8 hours dark cycles for 7 days. After 7 days, six of 12 inoculated and uninoculated seedlings from each variety were transferred to 1% agar plates containing 200 mM salt (NaCl) concentration while six were transferred to 1% agar plates without salt (control). Agar plates were transferred to the same growth chamber for 48 hours when seedlings were harvested. Shoots were collected for biomass measurements, roots were immediately frozen in liquid nitrogen, and stored in a −80°C freezer. Three seedlings per treatment per variety were used for WinRhizo scanning.

### Staining and visualization of roots using Light Microscopy

A modified method from Vierheilig *et al.* (1998) was used for staining roots. Details are provided in Supplemental Materials and Methods.

### GC-MS untargeted analysis for polar metabolites

Fifty (50 ± 2) mg (exact weight was recorded) of frozen root tissue was extracted, derivatised and analysed using GC-MS as described in Hill *et al.* (2013). Details are provided in Supplemental Materials and Methods.

### LC-MS untargeted analysis for lipid analysis

Lipids were extracted and analysed using LC-MS/MS using modified methods from Shiva et al. (2018) and Yu et al. (2018). Details are provided in Supplemental Materials and Methods.

### Data preprocessing and statistical analysis

Details for GC-MS and LC-MS data analysis and normalisation approaches are provided in Supplemental Materials and Methods. Data were log transformed and the Student’s *t*-test *p*-values determined. The *p*-values were further subjected to Benjamini Hochberg False Discovery Correction (Benjamini and Hochberg, 1995). The processed lipid data was subjected to multiple comparison statistical analyses using Analysis of Variance (ANOVA), with a false discovery rate (FDR)-adjusted *p* value of 0.05 and using the Benjamini–Hochberg method.

MetaboAnalyst 3.0 (Xia et al., 2015) software was employed for multivariate data analysis and additional *t*-test analysis. Fold-change for univariate analysis and bar plots were generated using GraphPad Prism 8.0 (GraphPad Software, Inc., USA).

## Supplemental Data

Supplemental Figure S1. WinRHIZO root scans of barley roots for cv. Vlamingh and cv. Gairdner genotypes. A- Vlamingh C, B- Vlamingh CF, C- Vlamingh S, D-Vlamingh SF; E- Gairdner C, F- Gairdner CF, G- Gairdner S, H- Gairdner SF. Abbreviations: Control- C, Control + fungus- CF, Salt- S and Salt + fungus- SF

Supplemental Figure S2. Principal component analysis (PCA) and heatmaps of four treatments in cv. Vlamingh (V) and cv. Gairdner (G). 2a- PCA of Control- C, Control + fungus- CF, Salt- S and Salt + fungus- SF in cv. Vlamingh; 2b- PCA of Control- C, Control + fungus- CF, Salt- S and Salt + fungus- SF in cv. Gairdner; 2c and 2d- heatmap obtained from hierarchical cluster analysis (HCA) of all identified metabolites in cv. Vlamingh and cv. Gairdner, respectively. The legend on the bottom indicates the sample groups and legends on the right are the identified metabolites.

Supplemental Figure S3. Metabolome and pathway view of alanine, aspartate and glutamate metabolism for metabolites with 2-fold change or greater in roots of barley cv. Vlamingh and cv. Gairdner under four treatments. 3a- Metabolome view showing all matched pathways arranged by p values on Y-axis and pathway impact values on X-axis. The node colour is based on its *p* value and the node radius is determined on their pathways impact values. 3b- Pathway view showing current metabolic pathway achieved by clicking the corresponding node on the metabolome view as given in 3a. Blue colour indicates default node and matched nodes are given in red colour. The compound names can be obtained via mouse over tool-tip.

Supplemental Figure S4. Heatmap showing the relative abundances of detected lipid species in samples from four treatments in two barley genotypes. Heatmap represents unsupervised hierarchical clustering of lipid species (rows). The row displays identified lipid species and the column represents the samples. Relative lower abundance of lipids in samples is displayed in blue, whilst relative higher abundance of lipids is displayed in red. Clustering was performed using Euclidean distance matrices on in the R package heatmap. VC- control treated cv. Vlamingh; VCF- control and fungus treated cv. Vlamingh, VSF- salt and fungus treated cv. Vlamingh; GC- control treated cv. Gairdner; GCF- control and fungus treated cv. Gairdner, GSF- salt and fungus treated cv. Gairdner. n = 6 for all treatments. salt- 200mM NaCl.

Supplemental Figure S5. Venn diagram summary of the number of significantly altered features identified by one-way ANOVA and post-hoc analysis of the lipid profiles from roots of two barley genotypes under salt stress and with and without fungal inoculation. The number represent significantly altered features regardless of their directionality (upregulation and downregulation). Numbers appearing in overlapped sections are common between treatments. Four analysis performed are Vlamingh salt (VS) compared to Vlamingh control (VC), Vlamingh salt and fungus (VSF) compared to Vlamingh salt (VS), Gairdner salt (GS) compared to Gairdner control (GC) and Gairdner salt and fungus (GSF) compared to Gairdner salt (GS).

Supplemental Figure S6. Flowchart for the experimental setup for plant growth conditions and inoculation with *Trichoderma harzianum* T-22.

Supplemental Tables S1. Identified metabolites from roots of two barley genotypes (cv. Vlamingh and cv. Gairdner) using gas chromatography- mass spectrometry. X-fold changes for treatment groups CF and S are compared against treatment C in both genotypes. X-fold changes for treatment SF is compared against treatment S in both genotypes. The identified metabolites are categorised in five categories- amino acids, organic acids, sugars, others and unknowns

Supplemental Tables S2. Average normalized responses of identified metabolites in roots of two barley genotypes- cv. Vlamingh and cv. Gairdner. Normalization was performed using fresh frozen weights of all samples and internal standard

Supplemental Tables S3. Detected m/z_RT mass features using MSDIAL from roots of two barley genotypes (cv. Vlamingh and cv. Gairdner). Four treatment groups were analysed- control, control + fungus, salt and salt+ fungus

Supplemental Tables S4. Annotated lipids returned with matches from MS-DIAL library in roots of two barley genotypes Four treatments were analysed for both genotypes- control, control + fungus, salt and salt + fungus

Supplemental Tables S5. Identified lipids with MS^2^ Spectra in roots of two barley genotypes Four treatments were analysed for both genotypes- control, control + fungus, salt and salt + fungus

Supplemental Tables S6. Fold changes of identified PCs- phosphatidylcholines and LPCs- lysophosphatidylcholines from roots harvested in agar study from four treatments- C-Control, S- Salt, CF- Control+Fungus, SF- Salt+Fungus, three comparisons were performed: S/C- salt vs control; CF/C- control+fungus/control; SF/S- salt+fungus/salt. V-Vlamingh, G-Gairdner

Supplemental Tables S7. Fold changes of identified PEs- phosphatidylethanolamines LPEs- lysophosphatidylethanolamines from roots harvested in agar study from four treatments- C- Control, S- Salt, CF- Control+Fungus, SF- Salt+Fungus, three comparisons were performed: S/C- salt vs control; CF/C- control+fungus/control; SF/S- salt+fungus/salt. V-Vlamingh, G-Gairdner

Supplemental Tables S8. Fold changes of identified DAG- diacylglycerol, TAG- triacylglycerol, MAG- monoacylglycerol, MGDG- monogalactosyl diacylglycerol, DGDG- digalactosyl diacylglycerol, ADGGA- Acyl diacylglyceryl glucuronide, from roots harvested in agar study from four treatments- C-Control, S- Salt, CF- Control+Fungus, SF- Salt+Fungus, Three comparisons were performed: S/C- salt vs control; CF/C- control+fungus/control; SF/S- salt+fungus/salt. V- Vlamingh, G-Gairdner

Supplemental Tables S9. Fold changes of identified DAG- diacylglycerol, TAG- triacylglycerol, MAG- monoacylglycerol, MGDG- monogalactosyl diacylglycerol, DGDG- digalactosyl diacylglycerol, ADGGA- Acyl diacylglyceryl glucuronide, from roots harvested in agar study from four treatments- C-Control, S- Salt, CF- Control+Fungus, SF- Salt+Fungus, Three comparisons were performed: S/C- salt vs control; CF/C- control+fungus/control; SF/S- salt+fungus/salt. V- Vlamingh, G-Gairdner

## Acknowledgements

The authors would like to thank A/Prof Stuart Roy (University of Adelaide) for providing barley seeds. We also thank Himasha Mendes for her help and support for GC-MS analysis. Lipid and metabolite analyses were performed at Metabolomics Australia at the University of Melbourne, which is a National Collaborative Research Infrastructure Strategy initiative under Bioplatforms Australia Pty Ltd (http://www.bioplatforms.com/). S.G gratefully acknowledges the David Lachlan Hay Memorial Fund (The University of Melbourne) for financial support during the preparation of this article.

